## Supplementary Information for "Capillary constrictions prime cancer cell tumorigenicity through PIEZO1"

### Supplementary Material

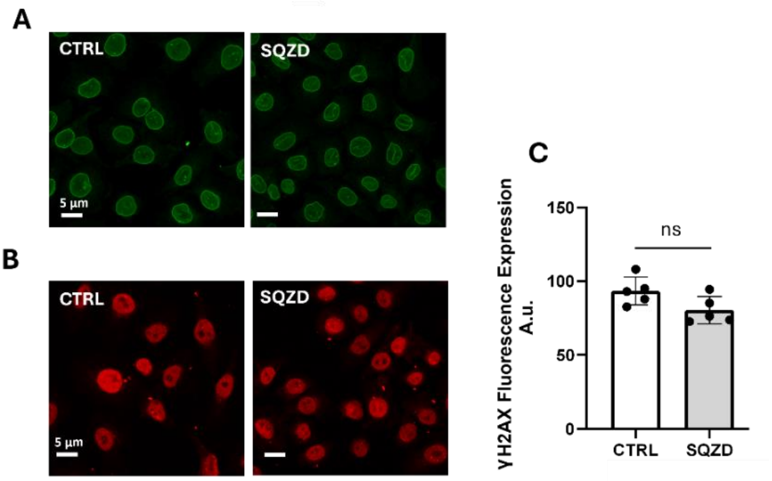


**Fig. S1 No significant differences in nuclear structure and DNA damage were observed between control and squeezed conditions. (A)** Representative staining of nuclear protein Lamin A in CTRL and SQZD melanoma cells. **(B)** Representative staining of γH2A.X in CTRL and SQZD melanoma cells. **(C)** Bar graph displaying the γH2A.x levels in CTRL and SQZD melanoma cells.

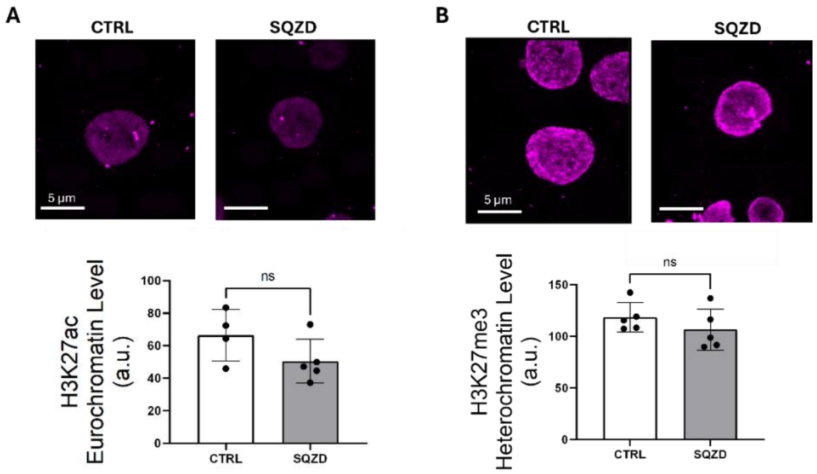


**Fig. S2 No significant differences in H3K27 acetylation or trimethylation were observed between control and squeezed conditions. (A)** Representative staining of H3K27ac in CTRL and SQZD melanoma cells. Bar graph displaying the H3K27ac levels in CTRL and SQZD cells. **(B)** Representative staining of H3K27m3 in CTRL and SQZD melanoma cells. Bar graph displaying the H3K9me3 levels in CTRL and SQZD cells.

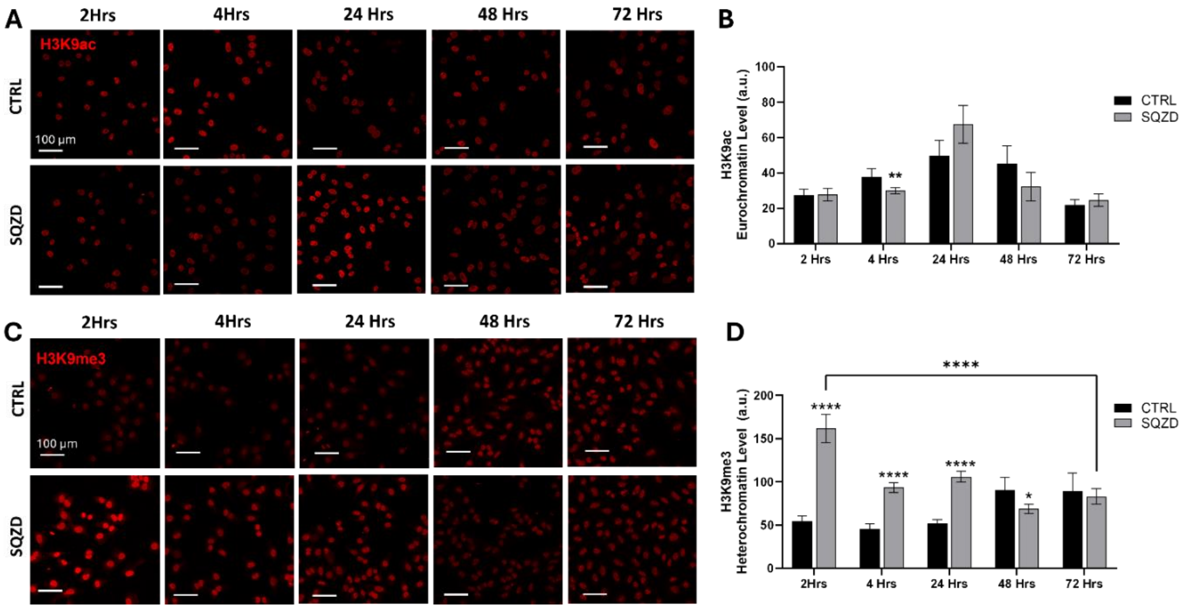


**Fig. S3. Temporal dynamics of histone modifications following mechanical deformation.** Representative immunofluorescence images of H3K9ac **(A)** and H3K9me3 **(C)** staining at different times post-constriction in control (CTRL) and squeezed (SQZD) A375-MA2 cells. Bar graphs displaying the quantification of ABCB5 H3K9ac **(B)** and H3K9me3 **(D)** mean fluorescence intensity levels in CTRL and SQZD cells at multiple time points (4 hrs to 72 hrs).

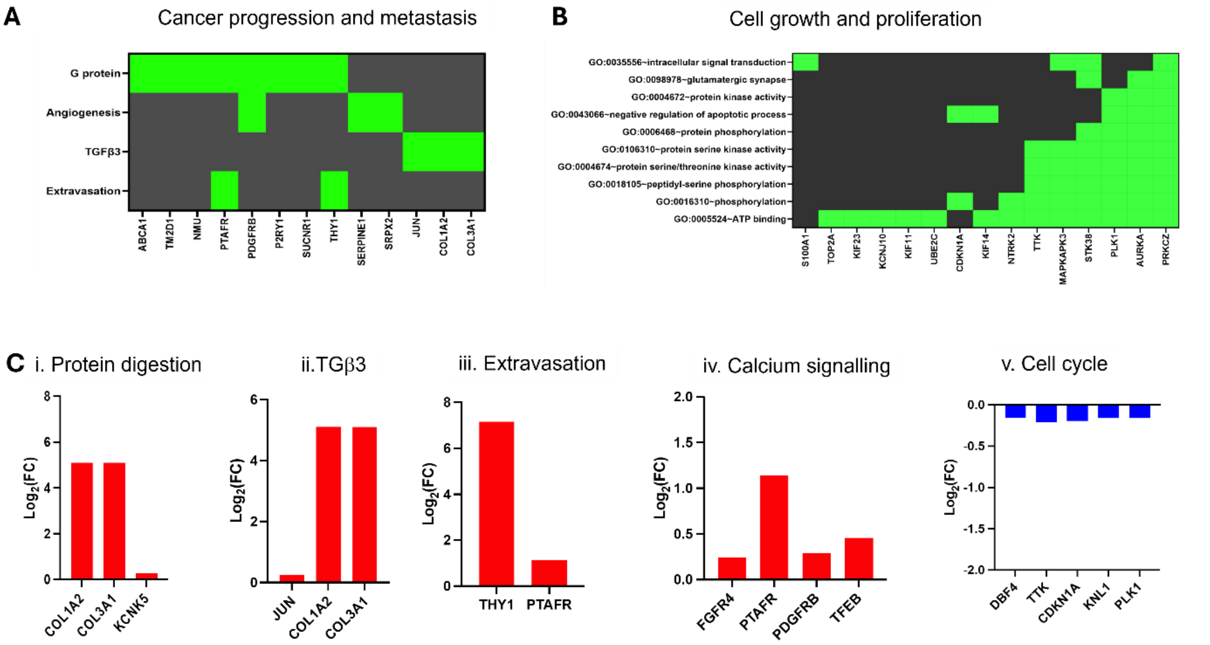


**Fig. S4 Gene ontology analysis. (A, B)** Enriched functional annotations for biological processes involved in cancer progression and metastasis and cell proliferation activity. **(C)** Graph charts representing genes involved in protein digestion (i), TGFβ3 pathway (ii), extravasation (iii), calcium signalling (iv) and cell cycle pathway (v).

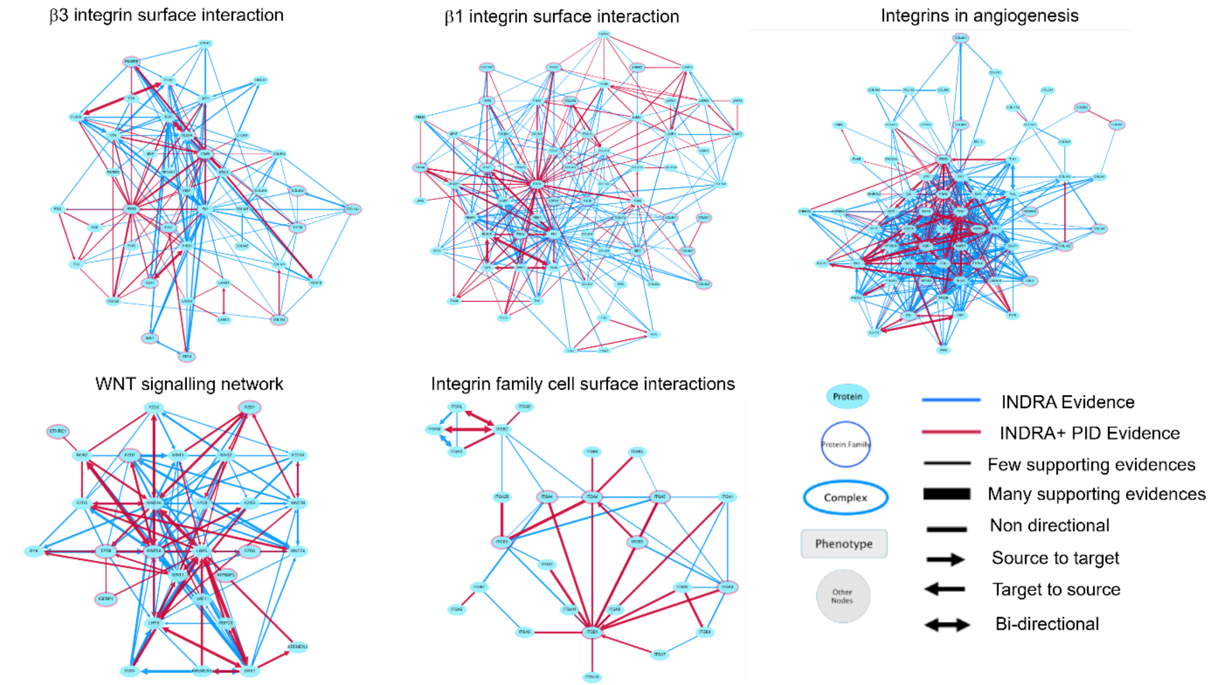


**Fig.S5 Analysis of gene function and construction of functional networks.** Gene networks representation derived from genes upregulated ($p<0.025$) fed to Cytoscape and analysed by KEGG database. Upregulated genes recognised in the different networks, were marked in red. INDRA: Integrated Network and Dynamical Reasoning Assembler; PID: Pathway interaction database.

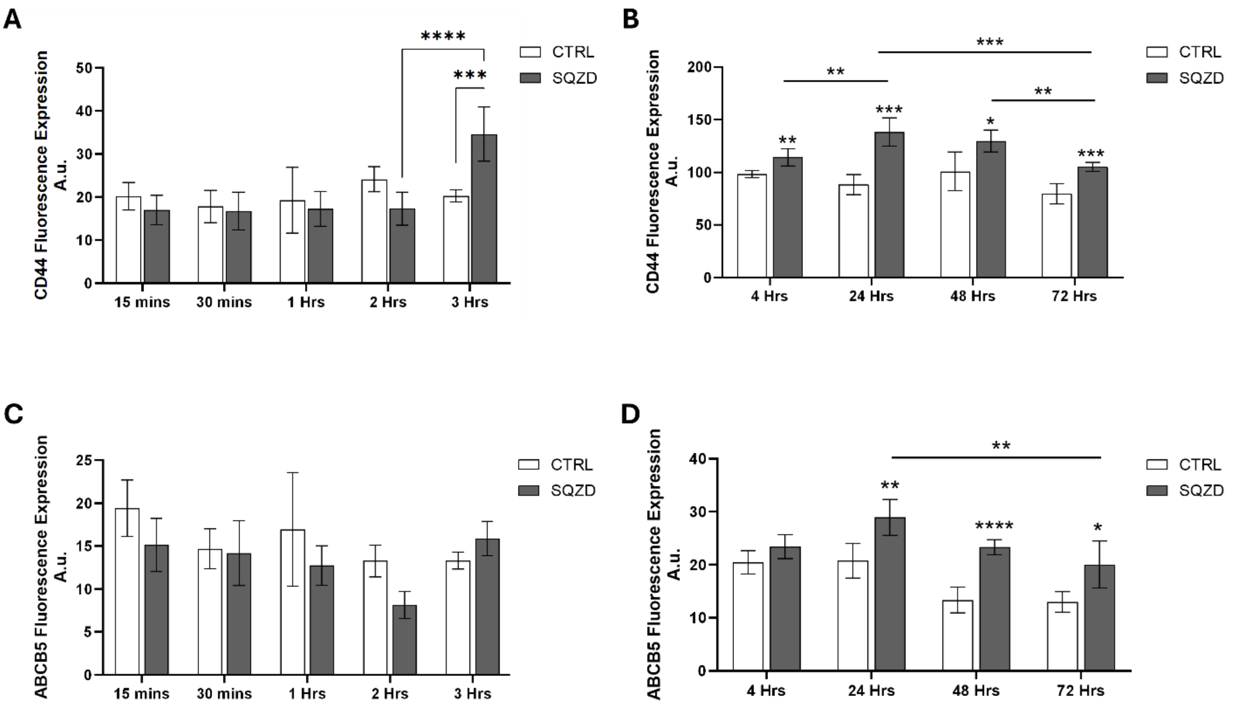


**Fig. S6 Temporal dynamics of phenotypic marker expression following transient mechanical constriction. (A)** Bar graph displaying the CD44 levels in CTRL and SQZD cells at multiple time points (15 min to 3 hrs). **(B)** Bar graphs displaying the CD44 levels in CTRL and SQZD cells at multiple time points (4 hrs to 72 hrs). **(C)** Bar graph displaying the ABCB5 levels in CTRL and SQZD cells at multiple time points (15 min to 3 hrs). **(D)** Bar graphs displaying the ABCB5 levels in CTRL and SQZD cells at multiple time points (4 hrs to 72 hrs).

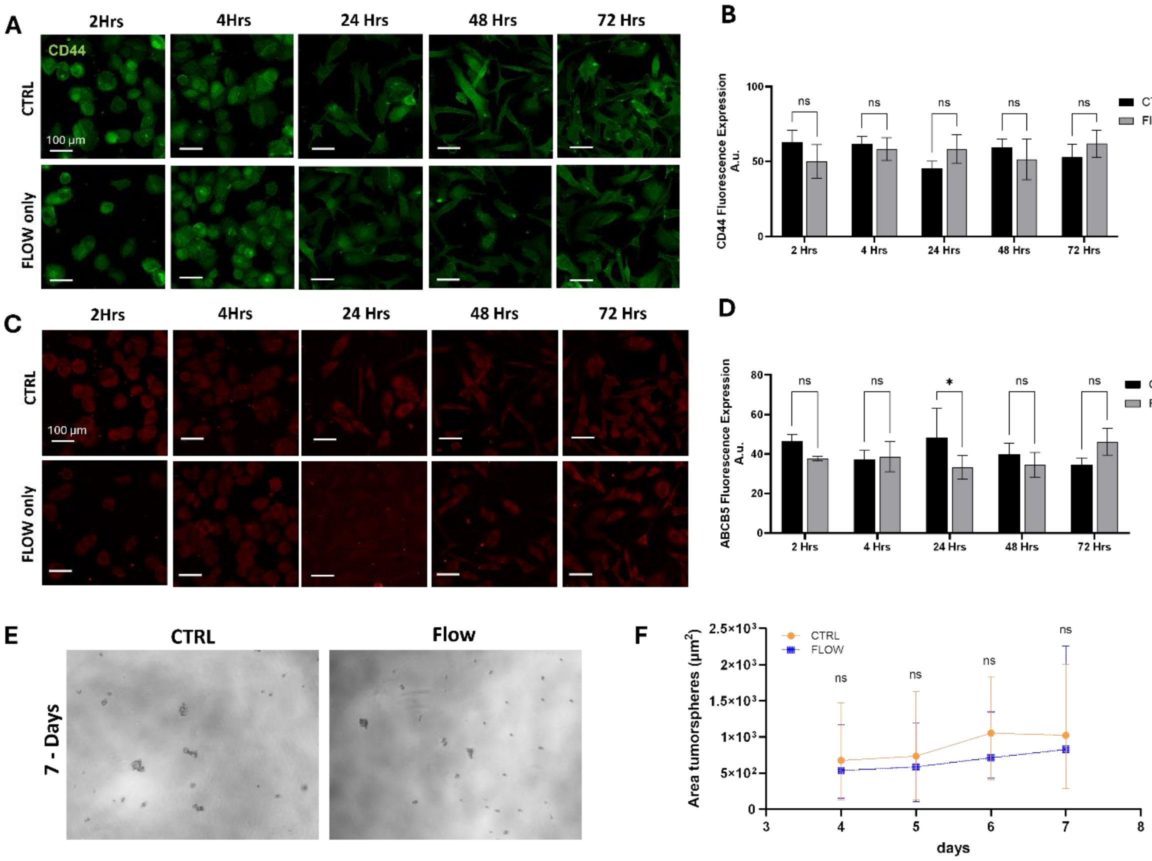


**Fig. S7 Mechanical deformation, not flow-induced shear stress, drives acquisition of stem-like traits.** Immunofluorescence analysis of CD44 (green, **A**) and ABCB5 (red, **C**) expression in control (static culture) and flow-only (no constriction) conditions. **(B, D)** Quantification of CD44 and ABCB5 fluorescence intensity. **(E)** Representative brightfield images of tumorsphere formed from control and flow-only groups. **(F)** Tumorsphere growth curves over time, showing no significant differences between control and flow-only conditions.

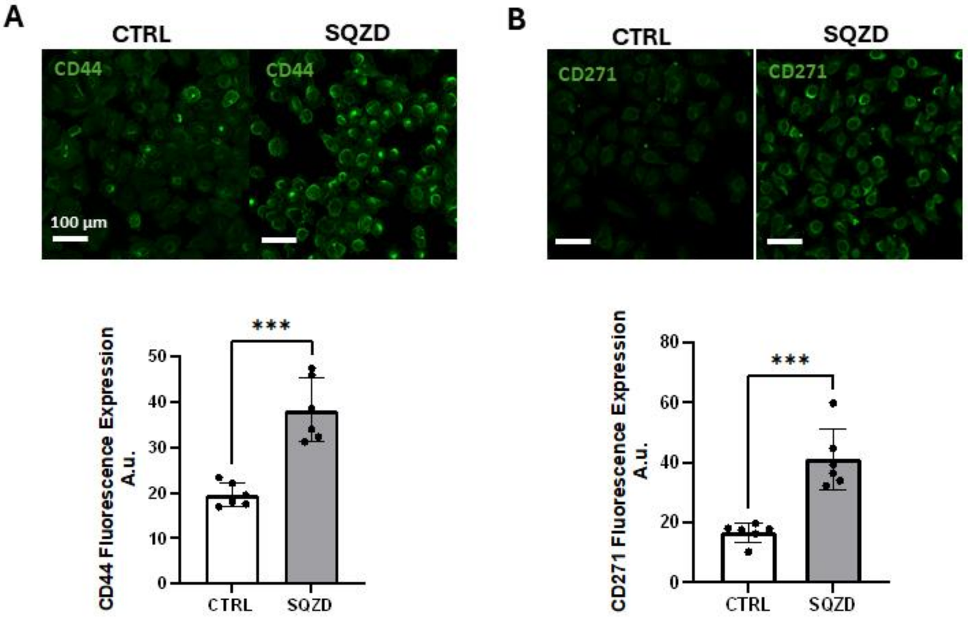


**Fig. S8 Verification of elevated tumorigenicity and stemness marker expression in A375-luciferase melanoma cells prior to in vivo experiments. (A)** Representative staining of CD44 in CTRL and SQZD melanoma cells. Bar graph displaying the CD44 levels in CTRL and SQZD cells. **(B)** Representative staining of CD271 in CTRL and SQZD melanoma cells. Bar graph displaying the CD271 levels in CTRL and SQZD cells.

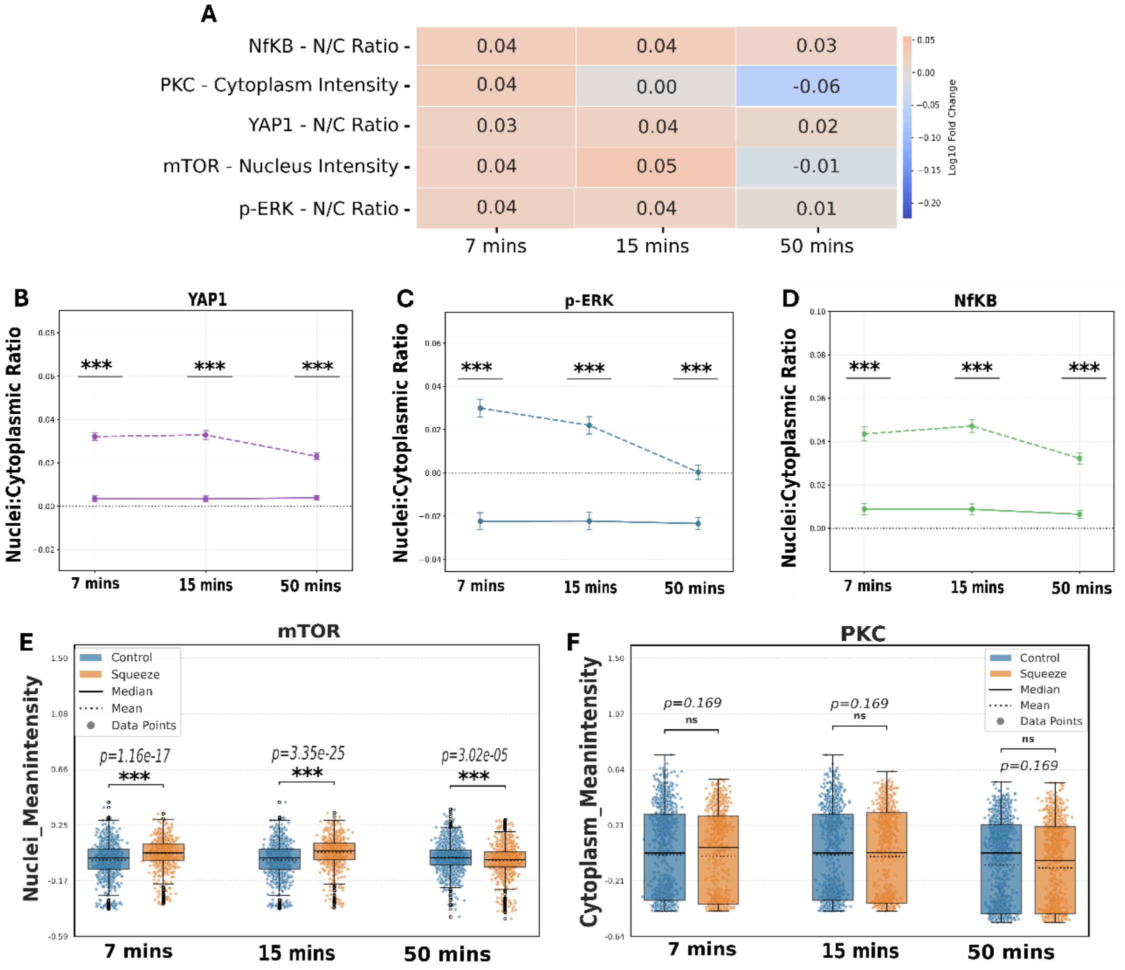


**Fig. S9. Phosphoprotein imaging reveals time-resolved activation of mechanosensitive signalling pathways. (A)** Heatmap representing $\log_{10}$ fold change in the activation of selected phospho-markers (NF-κB, PKC, YAP1, mTOR, and p-ERK) at 7-, 15-, and 50-minutes post-constriction. Line plot showing quantification of N/C ratios for YAP1 **(B)**, p-ERK **(C)**, and NF-κB **(D)** over time. Box plot representing mTOR nuclear intensity **(E)** and PKC cytoplasmic intensity **(F)** over time. Error bars indicate the Standard Error of the Mean (SEM).

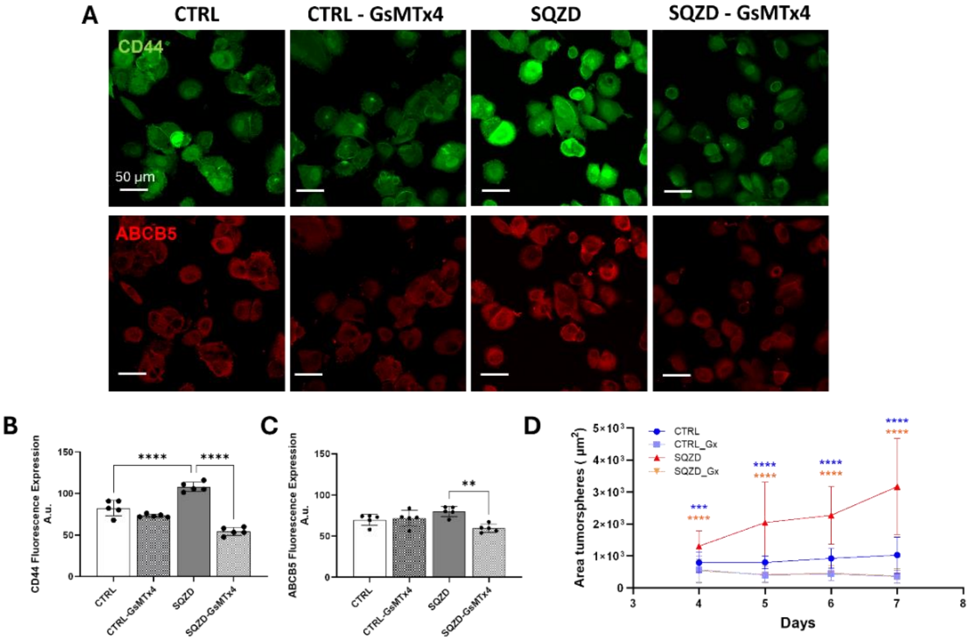


**Fig. S10 Inhibition of mechanosensitive channels with GsMTx4 reduces stemness marker expression and tumorsphere formation following mechanical squeezing. (A)** Representative immunofluorescence images of CD44 and ABCB5 expression in control (CTRL), squeezed (SQZD), and both cells group treated with GsMTx4. **(B, C)** Quantification of fluorescence intensity for CD44 and ABCB5 across experimental groups. **(D)** Graph representative of the mean tumorsphere area over time when cultured in serum-free media, for CTRL (blue), SQZD (red), and both cells group treated with GsMTx4 (light blue, orange).

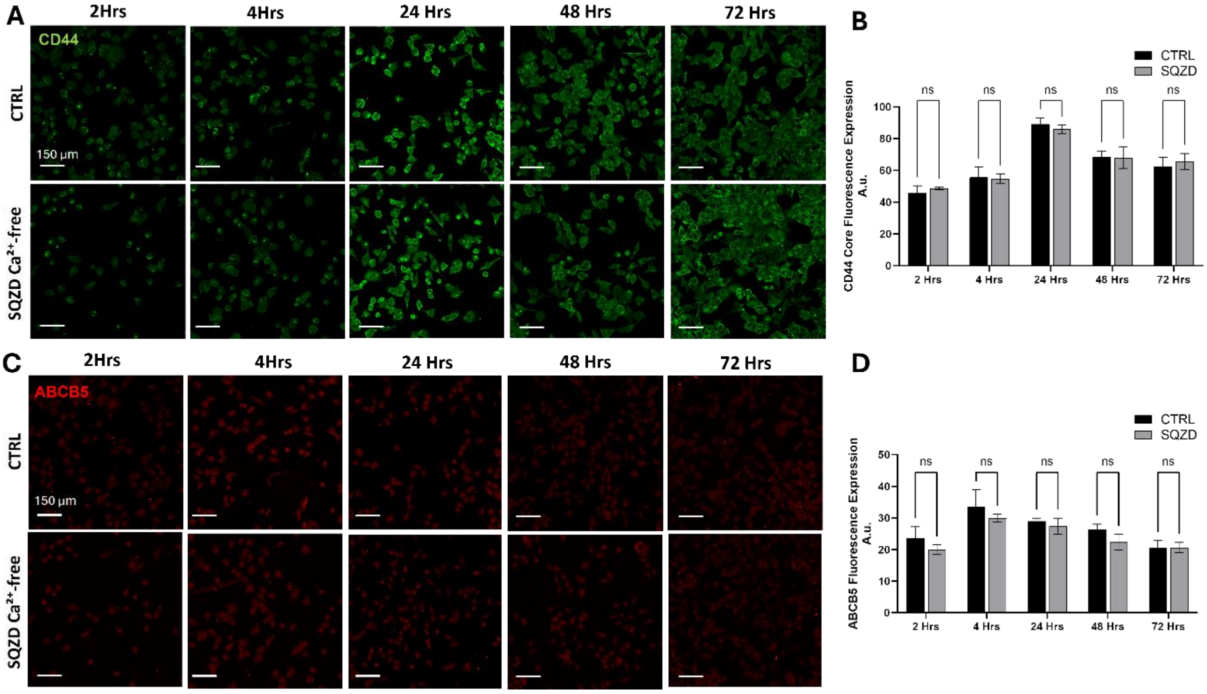


**Fig. S11 PIEZO1-induced stemness marker expression is dependent on extracellular calcium influx.** Representative immunofluorescence images showing CD44 **(A)** and ABCB5 **(C)** expression in melanoma cells under different conditions: control (CTRL) and squeezed cells incubated in calcium-free media during and after squeezing (SQZD $Ca^{2+}$-free). Quantification of fluorescence intensity for CD44 **(B)** and ABCB5 **(D)** across experimental groups.

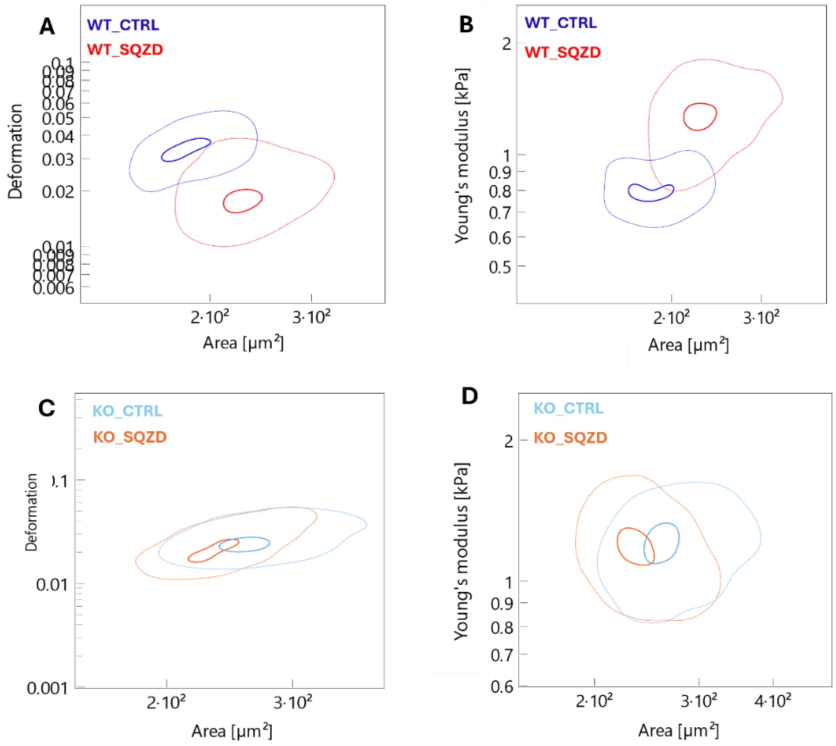


**Fig. S12 PIEZO1 is required for post-constriction mechanical adaptation in melanoma cells. (A–B)** Contour density plots showing the deformation **(A)** and apparent Young's modulus **(B)** of wild-type (WT) melanoma cells before (WT_CTRL, blue) and after (WT_SQZD, red) passing through a microfluidic constriction. **(C–D)** The same analysis for PIEZO1 knockout (KO) melanoma cells, comparing control (KO_CTRL, light blue) and constricted (KO_SQZD, orange) populations. Each contour plot represents the distribution of single-cell measurements (n > 1,000 cells per condition), with contour lines corresponding to 50% and 90% quantile density levels.

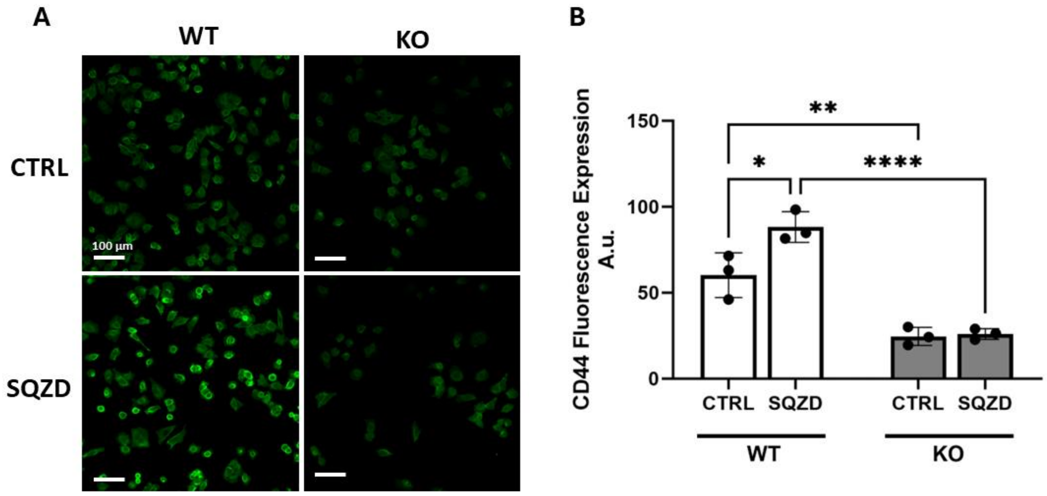


**Fig. S13 CD44 Expression in Cells Escaping from Transwell. (A)** Representative staining of CD44 in CTRL and SQZD melanoma cells (WT and KO). **(B)** Bar graph displaying the CD44 levels in CTRL and SQZD melanoma cells (WT and KO).

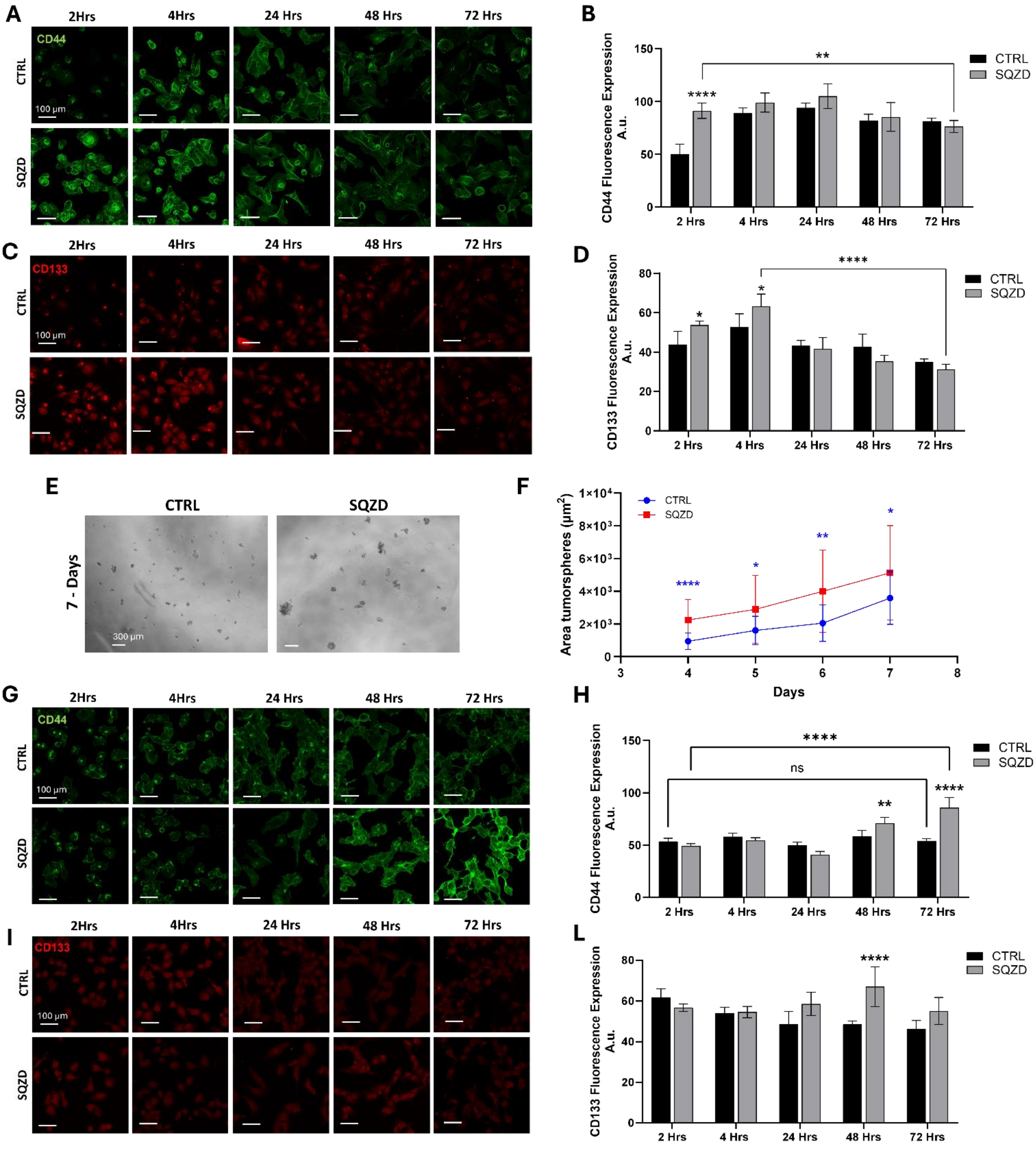


**Fig. S14 Mechanical constriction induces stemness markers and tumorsphere formation across additional cancer cell types.** Triple-negative breast cancer cells (MDA-MB-231) and osteosarcoma cells (HT-1080) were subjected to microcapillary constriction (SQZD) or maintained as controls (CTRL). Representative immunofluorescence images show expression of stemness markers CD44 (green) and CD133 (red) following mechanical constriction for MDA-MB-231 cells **(A, C)** and HT-1080 cells **(G, I).** Quantification of fluorescence intensity demonstrates upregulation of CD44 and CD133 expression after constriction in both cancer types; MDA-MB-231 cells **(B, D)** and HT-1080 cells **(H, L)**. **(E)** Representative brightfield images of tumorsphere formation for MDA-MB-231 cells. **(F)** Graph representative of the mean tumorsphere area over time when cultured in serum-free media, for CTRL (red), SQZD (red) MDA-MB-231 cells.

**Supplementary Table 1:** Indirect immunofluorescence labelling reagents

| Antibody | Dilution | Cat number | Host Species | Company |
|---|---|---|---|---|
| **p-ERK** | 1:200 | 9101S | Rabbit | Cell Signaling Technology |
| **p-AKT(308)** | 1:500 | 77440S | Rabbit | Cell Signaling Technology |
| **NFkB** | 1:200 | 8801S | Rabbit | Cell Signaling Technology |
| **p-p38** | 1:200 | 8632S | Rabbit | Cell Signaling Technology |
| **YAP1** | 1:500 | AB225440 | Rabbit | Abcam |
| **PKC** | 1:200 | MA1-157 | Mouse | Thermo Fisher Scientific |
| **mTOR** | 1:500 | 5048S | Rabbit | Cell Signaling Technology |
| **Anti-mouse AlexaFluor 647** | 1:1000 | 4410S | Goat | Cell Signalling Technology |
| **anti-rabbit AlexaFluor 555** | 1:1000 | 4413S | Goat | Cell Signalling Technology |
| **DAPI** | 1:1500 | D9542 | | Sigma-Aldrich |
| **Flash Phalloidin Green-488** | 1:100 | 424201 | | BioLegend |
